# Direct contribution of skeletal muscle mesenchymal progenitors to bone repair

**DOI:** 10.1101/2020.09.08.287367

**Authors:** Julien Anais, Kanagalingam Anuya, Megret Jérome, Luka Marine, Ménager Mickaël, Relaix Frédéric, Colnot Céline

## Abstract

Tissue regeneration relies on the activation of tissue resident stem cells concomitant with a transient fibrous tissue deposition to allow functional tissue recovery. Bone regeneration involves skeletal stem/progenitors from periosteum and bone marrow, the formation of a fibrous callus followed by the deposition of cartilage and bone to consolidate the fracture. Here, we show that mesenchymal progenitors residing in skeletal muscle adjacent to the bone fracture play a crucial role in mediating the initial fibrotic response to bone injury and also participate in cartilage and bone formation in the fracture callus. Combined lineage and scRNAseq analyses reveal that skeletal muscle mesenchymal progenitors adopt a fibrogenic fate before they engage in a chondrogenic fate after fracture. In polytrauma, where bone and skeletal muscle are injured, skeletal muscle mesenchymal progenitors fail to undergo fibrogenesis and chondrogenesis. This leads to impaired healing and persistent callus fibrosis originating from skeletal muscle. Thus, essential bone-muscle interactions govern bone regeneration through the direct contribution of skeletal muscle as a source of mesenchymal progenitors driving the fibrotic response and fibrotic remodeling, and supporting cartilage and bone formation.

## Introduction

Bone regeneration is usually described as a scarless and efficient regenerative process, beginning with an inflammatory response, the formation of a fibrous callus and the deposition of cartilage and bone tissues that are then slowly remodeled to reconstitute the initial shape and function of the injured bone. Skeletal/stem progenitors activated by the bone injury differentiate into chondrocytes preferentially in the center of the callus where endochondral ossification occurs, and the replacement of cartilage by bone is essential for successful healing. At the periphery of the callus where bone formation occurs via intramembranous ossification, skeletal/stem progenitors differentiate directly into osteoblasts. Several sources of skeletal stem/progenitors for bone repair have been identified including the bone marrow and the periosteum lying at the outer surface of bone ^1–5^. Other reports have pointed at the contribution of surrounding tissues such as skeletal muscle but the nature of the skeletal stem/progenitors from skeletal muscle and their role in bone regeneration remain undefined ^6–9^. The presence of bone forming cells in skeletal muscle has been suspected since Urist first showed that bone formation could be induced within skeletal muscle ^10^. The role of the myogenic lineage in bone repair has been previously investigated and revealed that the muscle stem cells, or satellite cells, were required for normal bone repair through their paracrine functions ^6^. Although satellite cells can differentiate into osteoblasts and chondrocytes *in vitro, in vivo* investigations suggested a poor osteochondrogenic potential of the myogenic lineage during bone repair ^6, 9, 11^. Here, we sought to explore other cell populations within the skeletal muscle interstitium, containing fibro/adipo progenitors (FAP) and mesenchymal progenitors (MP). These cell populations from the skeletal muscle interstitium are known to support skeletal muscle regeneration and become the source of persistent adipose and fibrotic tissue in pathological conditions such as muscular dystrophy ^12–16^ We characterized the skeletal muscle mesenchymal progenitors participating in cartilage and bone formation during bone repair and investigated their function in the context of musculoskeletal trauma. The role of intact skeletal muscle around bone is recognized clinically to be essential for bone repair as soft tissue damage can severely impair bone healing and skeletal muscle coverage can improve healing, but the underlying mechanisms are still unknown ^17–20^. Using a mouse polytrauma model and single-cell RNA-seq analyses of skeletal muscle mesenchymal progenitors, we uncover that the initial fibrotic response mediated by skeletal muscle mesenchymal progenitors and their commitment to the chondrogenic lineage in the fracture callus are impaired. Further, in the polytrauma environment skeletal muscle surrounding the bone fracture site is the source of persistent callus fibrosis compromising bone repair.

## Results

### Skeletal muscle contains a heterogeneous population of mesenchymal progenitors participating in bone repair

To elucidate the tissue origin of skeletal stem/progenitors cells, we co-transplanted Tomato-labelled EDL muscle (*Extensor Digitus Lengus*) and GFP-labelled periosteum grafts at the tibial fracture site of wild-type hosts. We observed concomitant recruitment of skeletal muscle and periosteal cells in cartilage and bone within the callus (Fig. 1a). We showed that the cellular contribution of skeletal muscle was physiological, by transplanting EDL muscle from *GFP-actin* mice adjacent to the tibia of a wild-type host and allowing complete regeneration of the transplanted EDL muscle for one month. Tibial fracture induced one-month post-EDL transplantation revealed similar contribution of skeletal muscle derived cells to cartilage and bone within the fracture callus (Fig. 1b).

**Figure 1:**
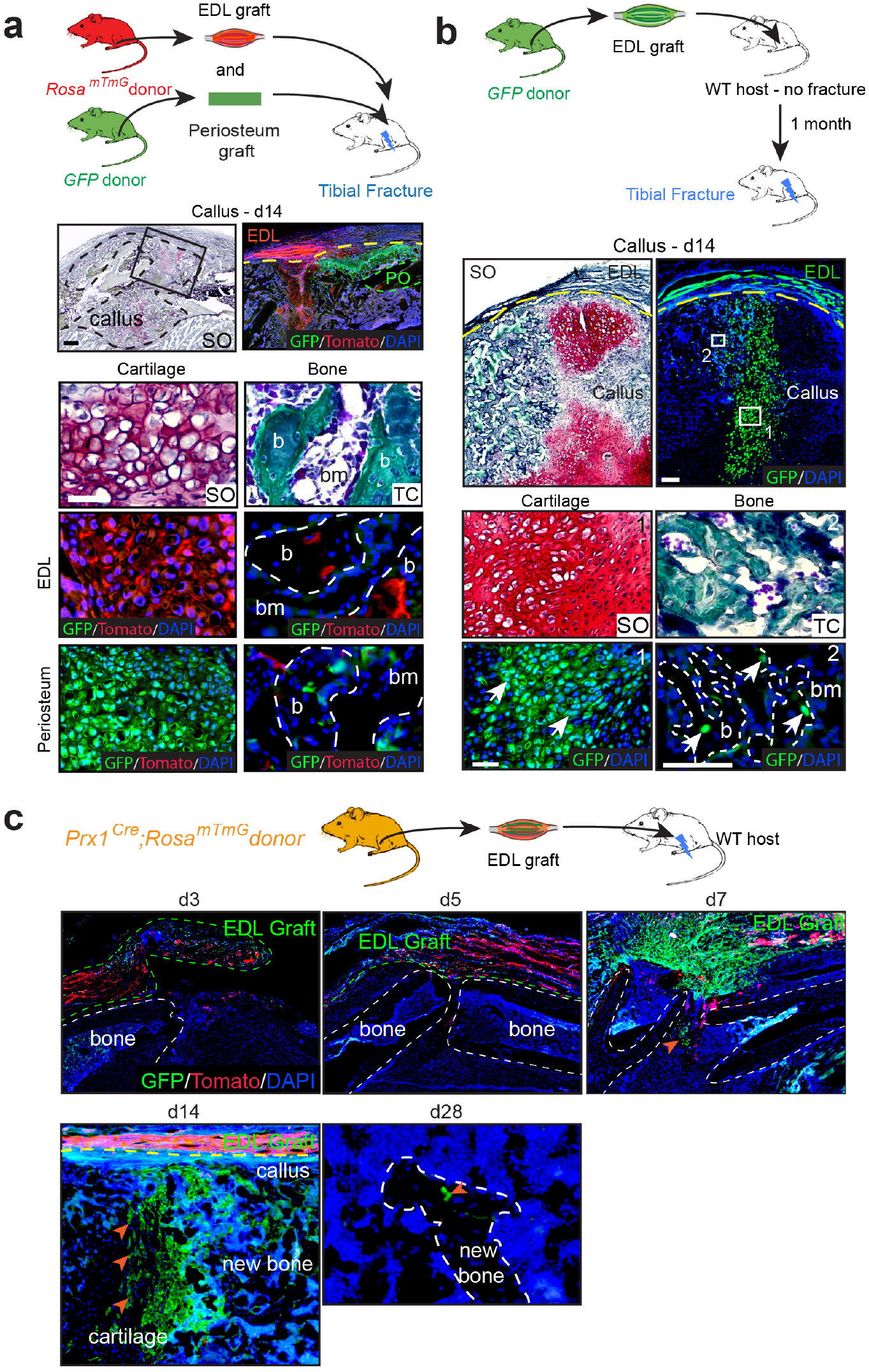
Skeletal-muscle is a source of osteochondroprogenitors during bone repair. **a**, Combined grafting of periosteum from *GFP-actin* mice and EDL-muscle from *mTmG* mice at the fracture site of wild-type hosts. Longitudinal sections of tibial callus (delimited by a black dotted line) at 14 days post-fracture and stained with Safranin-O (SO, left) and enlarged view of boxed region on adjacent section counterstained with DAPI (right). EDL muscle graft outside the callus and skeletal muscle-derived cells within the callus (Tomato+ signal) (callus limited by a yellow dotted line). Periosteum graft (PO, delimited by a green dotted line) and periosteum-derived cells within the callus (GFP+ signal). High magnifications of cartilage and bone (b) derived from the EDL muscle graft ^31^ or from the periosteum graft (green) stained with SO and Masson’s Trichrome (TC) and adjacent sections counterstained with DAPI (bone delimited by a white dotted line). b, Top: Experimental design of tibial fracture induced one month after GFP-EDL muscle graft transplantation. Middle: Callus sections of GFP-EDL muscle graft next to the fracture callus (delimited by a yellow dotted line) at d14 post-fracture stained with SO (left) and counterstained with DAPI (right). Bottom: High magnification of cartilage (box 1) and bone (box 2, white dotted line) containing GFP+ EDL muscle-derived chondrocytes and osteocytes respectively (white arrow). c, Experimental design of *Prx1^Cre^;Rosa^mTmG^* EDL muscle graft transplanted at the fracture site of wild-type hosts. Representative sections of fracture calluses at days 3, 5, 7, 14 and 28 post-fracture showing skeletal muscle derived cells in callus (orange arrowhead). Scale bar: SO/TC=1mm, high magnification= 50μm for cartilage and 25 μm for bone, bm: bone marrow. Representative images of 3 distinct samples.

To characterize the cartilage and bone forming cells derived from skeletal muscle, we induced a fracture in *Pax7C^reERT2^;Rosa^mTmG^* mice or transplanted EDL-skeletal muscle grafts from *Pax7^CreERT2^;Rosa^mTmG^* mice at the fracture site of wild-type hosts. Tomato-positive cells were detected within callus but no GFP-positive cells, demonstrating the absence of contribution from the myogenic lineage (Extended Data Fig. 1a, b). All skeletal stem/progenitors forming cartilage and bone in the fracture callus are derived from the Prx1 mesenchymal lineage that marks bone marrow stromal/stem cells and periosteal cells ^3^. Transplantation of EDL grafts from *Prx1^Cre^;Rosa^mTmG^* donors showed that transplanted skeletal muscle also contains osteochondroprogenitors for bone repair strictly derived from the Prx1 mesenchymal lineage. These osteochondroprogenitors started migrating at the center of the callus from skeletal muscle adjacent to bone between days 5 and 7 post-fracture and were detected within cartilage by day 7 and within bone until day 28 (Fig. 1c). To confirm the presence of osteochondroprogenitors within skeletal muscle, we showed that mononucleated cells isolated from *Prx1^Cre^;Rosa^mTmG^* hindlimb muscles free of fascia, tendon and fat, and transplanted in wild-type host, were able to integrate into callus cartilage and bone (Extended Data Fig. 1c). At steady state, on transverse sections of *Prx1^Cre^;Rosa^mTmG^* TA muscle, the Prx1-derived GFP-labelled cells localized in the skeletal muscle interstitium next to capillaries, co-expressed the pericyte markers, NG2 and PDGFRβ, and the mesenchymal markers, PDGFRα and CD29 (Fig. 2a). To better understand the cellular composition of the skeletal muscle mesenchymal progenitor population, we performed scRNA-seq analysis of sorted Prx1-derived skeletal muscle cells surrounding the tibia. We identified 9 clusters and defined four sub-populations, distinct from endothelial and hematopoietic cell populations, and including FAP/MP (expressing *Prrx1, Cxcl12, Pdgfra, Ly6a* and *Cd34*), tenocyte-like cells (expressing *Scx, Tnmd* and *Kera*), pericytes (expressing *Cspg4, Des* and *Mylk*) and *Spp1+/Lgals3+* cells (Figure 1b-e and Extended Data Fig. 2a). Additionally, flow cytometry analyses showed that freshly isolated Prx1-derived skeletal muscle cells represent 33,5% of mononucleated cells within muscle, are CD45-CD11b-CD31- and coincide with populations expressing the pericyte/mesenchymal marker PDGFRβ, the fibroadipoprogenitor/mesenchymal progenitor (FAP/MP) markers CD29, PDGFRα, Sca1 and CD34, but do not encompass all mesenchymal cell types within skeletal muscle (Extended Data Fig. 2b-d). Cultured GFP+ Prx1-derived skeletal muscle cells exhibited osteogenic, adipogenic, chondrogenic and fibrogenic potential but no myogenic potential and expressed fibro-mesenchymal, pericyte and tenocyte markers (Extended Data Fig. 2e, f). Skeletal muscle thus contains a heterogeneous population of skeletal muscle mesenchymal progenitors derived from Prx1 mesenchymal lineage and contributing to bone repair.

**Figure 2:**
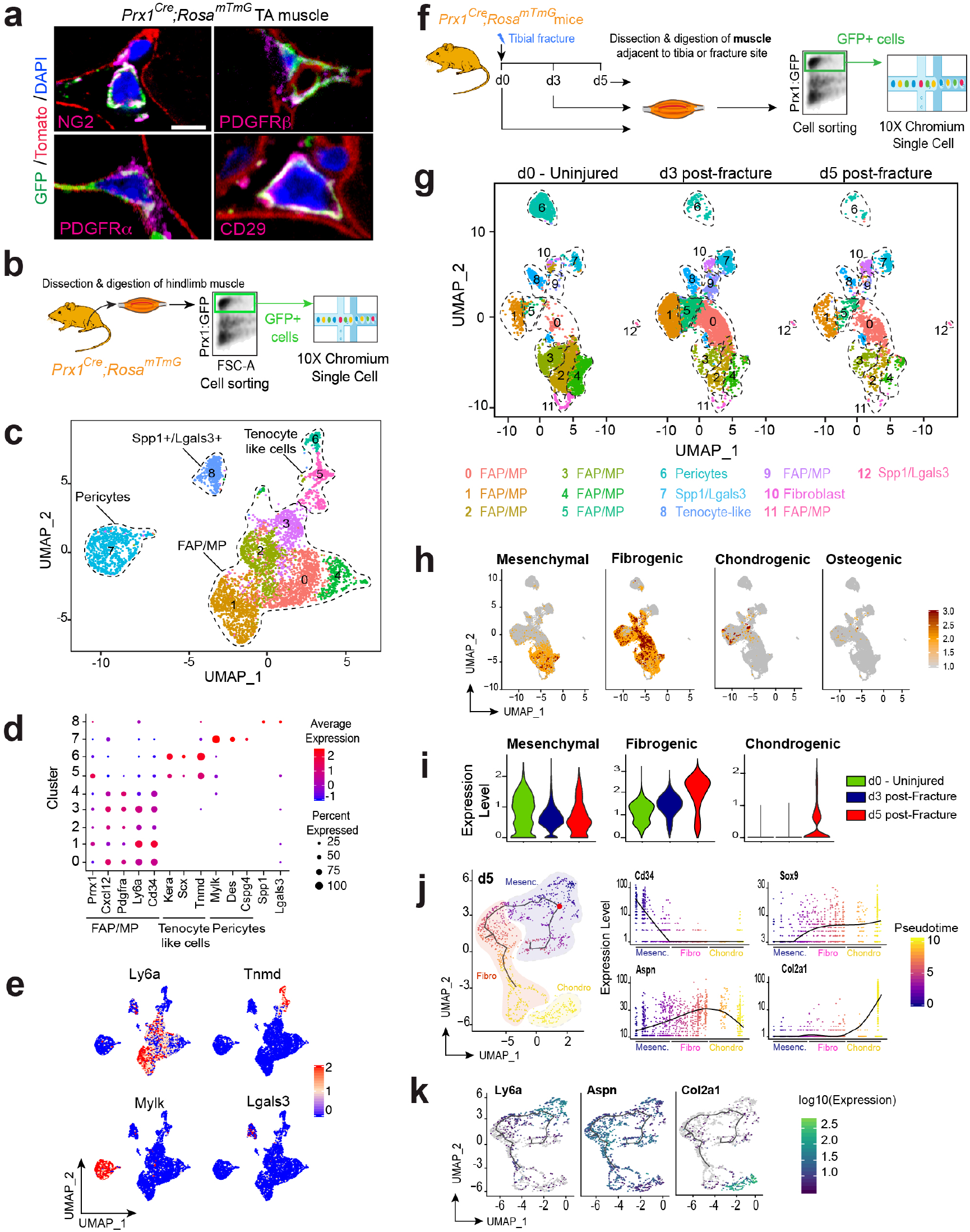
Single cell RNAseq analyses of skeletal muscle mesenchymal progenitors in intact muscle and in response to fracture. **a**, Transverse sections of TA muscle from *Prx1^Cre^;Rosd^mTmG^* mice immunostained with NG2, PDGFRβ, PDGFRα or CD29 antibodies (magenta). DAPI labelled nuclei, GFP labelled Prx1-derived cells and Tomato labelled non Prx1-derived cells. Representative images of 3 distinct samples. b, Experimental design of scRNAseq experiment. Prx1-derived skeletal muscle cells were isolated from skeletal muscle and sorted based on GFP-expression prior to scRNA-seq. c, UMAP projection of color-coded clustering of Prx1-derived cells reveals 9 clusters defining 4 distinct populations (limited by a black dotted line). d, Dotplot of indicated genes expression identifying FAP/MP, tenocyte-like cells, pericytes and Spp1/Lgals3+ cell populations. e, FeaturePlot of sub-population marker expression (*Ly6a* for FAP/MP, *Tnmd* for tenocyte like cells, *Mylk* for pericytes and Lgals3 for Spp1+/Lgals3+ cells). f, Experimental design of single-cell analyses. Prx1-derived skeletal muscle cells were isolated from skeletal muscle at day 0, day 3 post-fracture and day 5 post-fracture and sorted based on GFP-expression prior to scRNA-seq. The 3 expression matrices were integrated to generate the dataset used for the analysis. g, UMAP projection of color-coded clustering of combined d0, d3 and d5 samples. Cluster identities are indicated below. h, FeaturePlot of mesenchymal, fibrogenic, chondrogenic and osteogenic lineages scores. i, Pseudobulk expression of mesenchymal, fibrogenic and chondrogenic score in d0, d3 and d5 post-fracture samples. j, Pseudotime analysis of FAP/MP at d5 post-fracture (left). Expression of *Ly6a, Aspn, Sox9* and *Col2a1* genes along pseudotime (right). k, FeaturePlot of *Ly6a, Aspn* and *Col2a1* expression as example of mesenchymal, fibrogenic, chondrogenic lineages in d5 post-fracture sample. Scale bars: a= 10μm.

### Skeletal muscle mesenchymal progenitors undergo fibrogenesis before chondrogenesis in response to bone fracture

We then characterized the cellular response of skeletal muscle mesenchymal progenitors to bone fracture. Skeletal muscle surrounding the fracture site was dissected, excluding periosteum and bone marrow tissues. After enzymatic digestion, we sorted Prx1-derived GFP+ cells and performed scRNA-seq analysis combining d0 (un-injured), d3 and d5 post-fracture samples in a common dataset (Figure 2F). We identified 13 clusters in skeletal muscle mesenchymal progenitors that can be partitioned into 4 distinct populations (FAP/MP, tenocyte-like cells, pericytes, and *Spp1*/Lgals3 cells) already defined in un-injured skeletal muscle and a distinct fibroblast cluster (Fig. 2f-g, Extended Data Fig. 3a, b). This fibroblast cluster was defined as cells expressing genes coding for ECM proteins (*Col1a1, Sparc, Col3a1, Col5a1* and *Postn*) but no other subpopulation markers (Extended Data Fig. 3c). To assess the fate of skeletal muscle mesenchymal progenitors in response to fracture, we annotated the dataset according to known lineage markers: mesenchymal (MP), fibrogenic (ECM producing cells), chondrogenic and osteogenic (Table1). Mesenchymal and fibrogenic lineage markers were expressed mostly by FAP/MP at d0, d3 and d5 while chondrogenic lineage markers were only expressed at d5. Osteogenic lineage markers were expressed at low level in skeletal muscle mesenchymal progenitors at d3 and d5 (Fig. 2h, i and Extended Data Fig. 2d). *In silico* trajectory analysis on d5 post-fracture sample showed that skeletal muscle mesenchymal progenitors express mesenchymal lineage markers and start expressing fibrogenic lineage markers in response to fracture prior to chondrogenic markers, except for Sox9 which is already detected in the fibrogenic state (Fig. 2j, k). These results indicate that skeletal muscle mesenchymal progenitors upon activation engage in a fibrogenic fate before undergoing early chondrogenic differentiation from d5 post-fracture. This cellular response to injury occurs specifically within the FAP/MP population of skeletal muscle mesenchymal progenitors.

### Musculoskeletal trauma impairs bone healing

To determine the role of skeletal muscle mesenchymal progenitors in musculoskeletal trauma, where concomitant injury of bone and muscle compromises skeletal regeneration, we developed a clinically relevant polytrauma mouse model. As observed in human, mechanical injury to skeletal muscles surrounding a fractured tibia caused fracture non-union, displayed by (*i*) delayed in callus, cartilage and bone formation by day 7, (*ii*) impaired cartilage and bone resorption, (*iii*) abnormal fibrous tissue accumulation, and (*iv*) absence of bone bridging through day 56 (Fig. 3a, b and Extended Data Fig. 4a). This fracture non-union phenotype was correlated with decreased contribution to cartilage of skeletal muscle cells (Fig. 3c, d). The mechanical injury to skeletal muscle alone led to heterogeneous and delayed muscle regeneration as shown by areas containing regenerating muscle fibers and areas containing fibrous tissue at days 14 and 30 post-injury (Extended Data Fig. 4b). Bone healing was not impaired when combined with TA muscle injury only, thus a threshold of soft tissue trauma exists above which bone healing cannot occur efficiently (Extended Data Fig. 4c).

**Figure 3:**
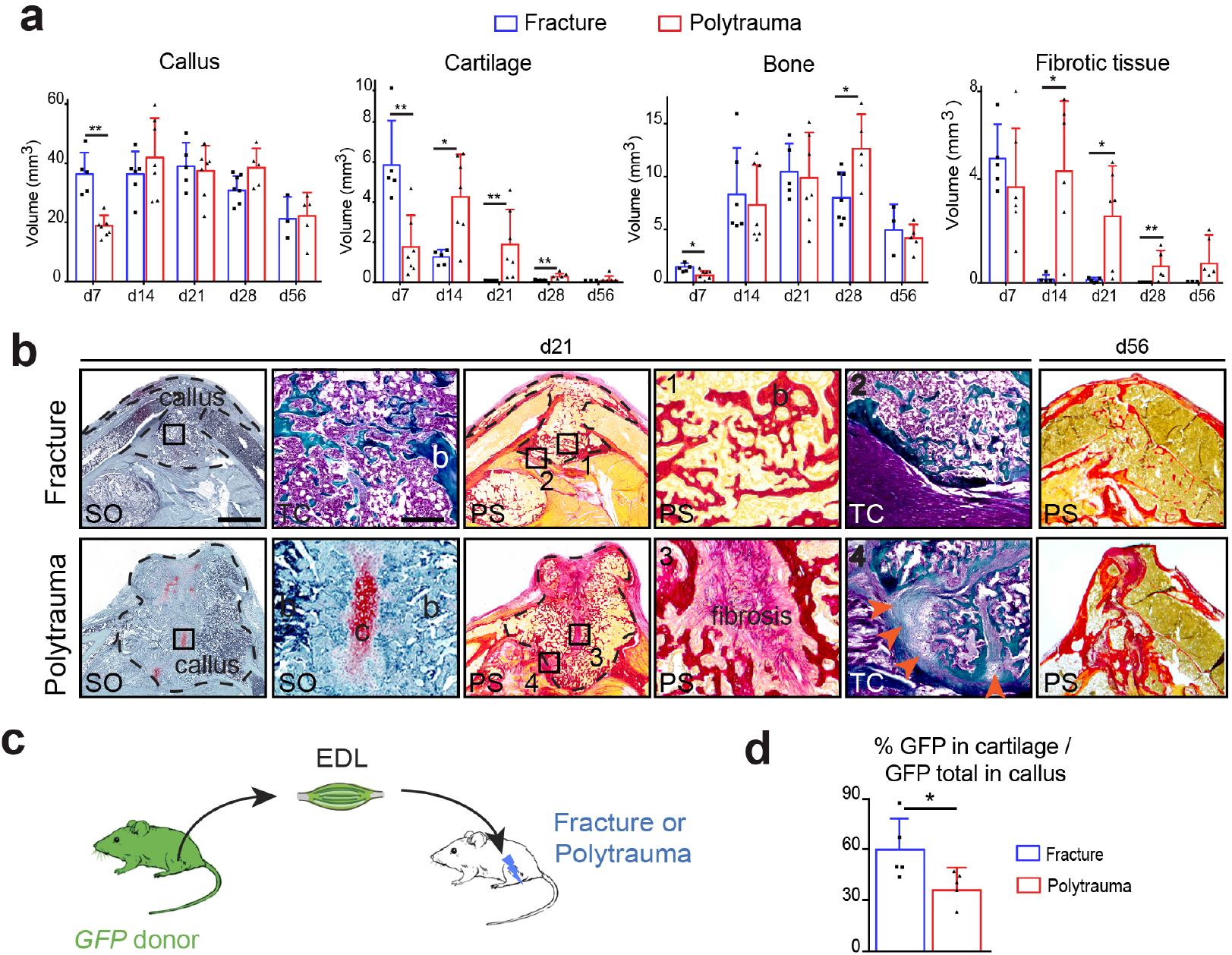
Polytrauma severely impairs bone healing and cellular recruitment from skeletal muscle. **a**, Histomorphometric quantification of callus, cartilage, bone and fibrotic tissue volumes in tibial fracture or in polytrauma through all stages of bone repair (d7 n=5 and n=7; d14, n=5 and n=7, d21 n=5 and n=7, d28 n=7 and n=5, d56 n=3 and n=5 for fracture and for polytrauma respectively). **b**, Representative callus sections stained with Safranin-O (SO), Trichrome (TC) and PicroSirius (PS) at days 21 and 56 (b: bone). Fully ossified callus and bone bridging in fracture (boxes 1, 2). Unresorbed cartilage (c), fibrosis (box 3) and absence of bone bridging (box 4, orange arrowheads) in polytrauma. Scale bars: low magnification = 1mm; boxed areas=200μm. **c**, Experimental design of EDL muscle graft from *GFP-actin* mice transplanted at the fracture site of tibial fracture or polytrauma. **d**, Histomorphometric analyses of GFP+ cartilage at day 14 post-injury. n=5 per group. Statistics: two-tailed Mann-Whitney test, * *P*<0,05; ** *P*<0,01, all data represent mean ± SD.

### Musculoskeletal trauma alters early fibrotic response of skeletal muscle mesenchymal progenitors to fracture

To understand how polytraumatic injury impacts skeletal muscle mesenchymal progenitors activation and differentiation, we performed scRNAseq analyses of skeletal muscle mesenchymal progenitors after fracture or polytrauma (Fig. 4a). Analysis of combined d0, d3 post-fracture, d5 post-fracture, d3 post-polytrauma and d5 post-poly-trauma samples showed that d0 sample was distinct to others samples. Samples from d3 post-fracture and d3 post-polytrauma clustered together separately from d5 post-fracture and d5 post-polytrauma samples that also clustered together (Fig. 4b left). Combined analysis of the 5 experimental groups uncovered 17 clusters highlighting 6 populations: FAP/MP, fibroblasts, tenocyte-like cells, pericytes, Spp1+/Lgals3+ cells and chondrocytes. Chondrocytes were defined as cells expressing *Col2a1, Acan* and *Fgfr3* (Fig. 4b right, c and Extended Data Fig. 5a, b). Analysis of cell cycle showed no difference in cell proliferation between fracture and polytrauma (Fig. 4d and Extended Data Fig. 5c). We then determined whether polytrauma has an impact on the capacity of skeletal muscle mesenchymal progenitors to engage into fibrogenic and chondrogenic fate compared to fracture. Analyses of mesenchymal and fibrogenic marker expression suggested a delay in down-regulation of mesenchymal markers at d3 post-polytrauma, correlated with a delay in up-regulation of fibrogenic markers at d3 and d5 post-polytrauma. In polytrauma, the chondrogenic markers were not expressed at d5 compared to fracture (Fig. 4e). We further focused gene ontology (GO) analysis on non-proliferative clusters 2 and 3 for d3 analysis and cluster 8 for d5 analysis, according to the percentage of cells from each sample per cluster and the cell cycle state (Fig. 4d, f). We used differentially expressed genes between clusters 3 (composed of d3 post-fracture cells) and 2 (composed on d3 post-polytrauma cells), and between post-fracture and post-polytrauma cells within cluster 8 to run GO analysis. In response to fracture at d3, cells have a high metabolic activity and secrete ECM. In response to polytrauma, cells express markers of angiogenesis, hypoxia and immune response but exhibit a lower metabolic activity and are not secreting ECM (Fig. 4g). At d5 post-fracture, cells expressed high level of ECM genes, genes from TGFβ pathway and signalling pathways linked with chondrogenic differentiation (Table2). These changes were not observed in d5 post-polytrauma. Instead cells exhibited higher level of metabolic activity suggesting a delay in their activation post-injury (Fig. 4h). These results show that the fibrogenic engagement of skeletal muscle mesenchymal progenitors is a crucial step during bone repair and precedes chondrogenic differentiation. In polytrauma, skeletal muscle injury perturbs the commitment toward the fibrogenic fate and delays the chondrogenic differentiation of skeletal muscle mesenchymal progenitors.

**Figure 4:**
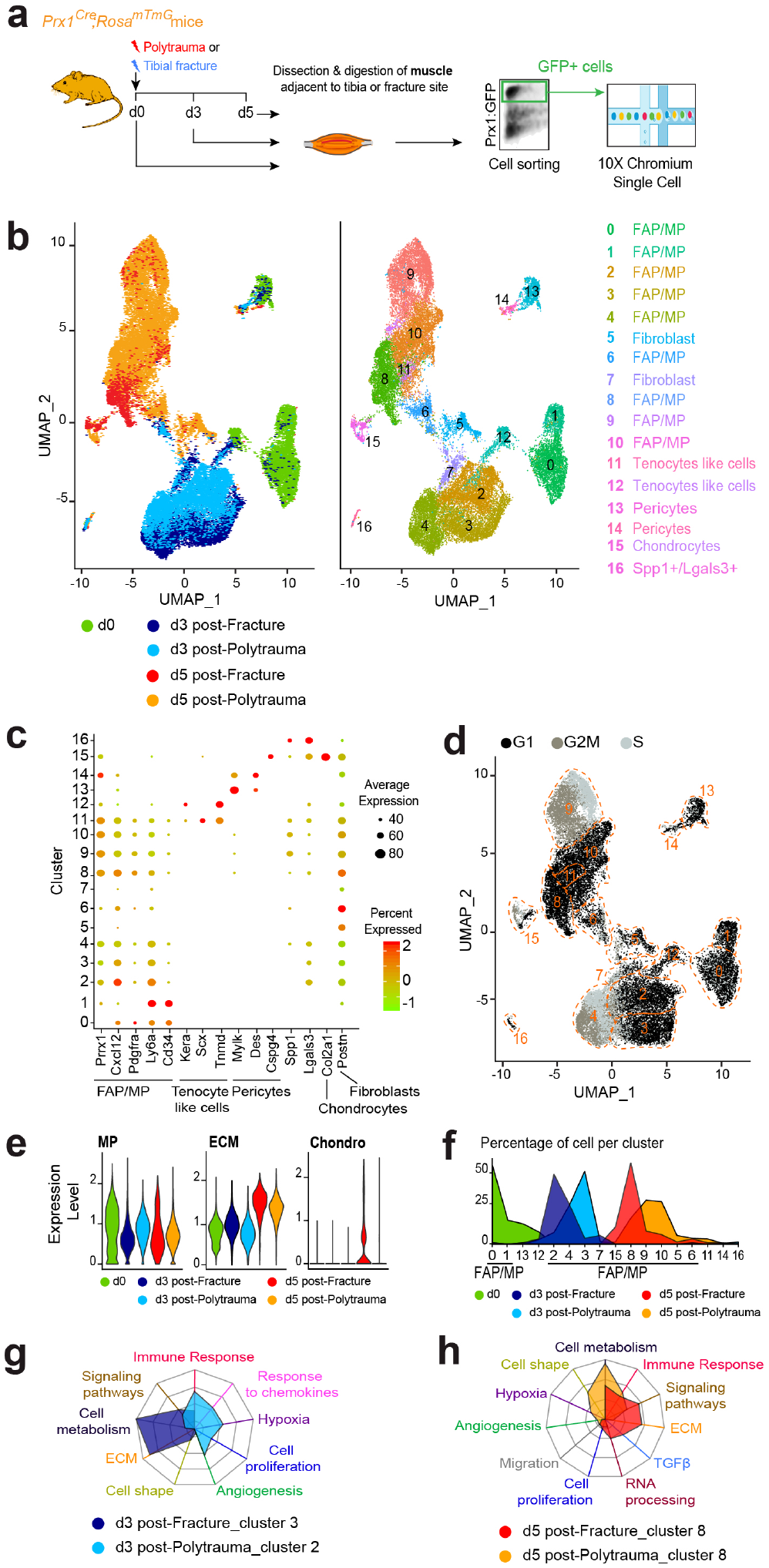
Single-cell analyses of skeletal muscle mesenchymal progenitors reveal impairment of initial fibrotic response in polytrauma. **a**, Experimental design of scRNAseq experiment. Prx1-derived skeletal muscle cells were isolated at d0, d3 and d5 post-fracture or post-polytrauma and sorted based on GFP-expression prior to scRNA-seq. The 5 expression matrices were combined to generate the dataset used for the analysis. b, UMAP projection of color-coded sample (left) and clustering (right) of combined analysis of d0, d3 and d5 post-fracture and post-polytrauma samples. Sample and cluster identities are indicated to the bottom and the right respectively. c, Dotplot of indicated genes expression identifying FAP/MP, tenocyte-like cells, pericyte, Spp1+/Lgals3+, chondrocyte and fibroblast cell populations. d, UMAP representation of color-coded cell cycle phases. e, Pseudobulk expression of mesenchymal, fibrogenic and chondrogenic score in d0, d3 and d5 post-fracture and post-polytrauma samples. f, Percentage of cell per cluster according to sample. g, Radar chart of enriched Gene Ontology functions in cluster 3 corresponding to d3 post-facture and cluster 2 corresponding post-polytrauma. h, Radar chart of enriched Gene Ontology functions in d5 post-fracture *versus* post-polytrauma in cluster 8.

### Skeletal muscle is the source of persistent callus fibrosis in traumatic injuries

We then analyzed the consequences of polytrauma at later stages of repair. As observed by scRNAseq analysis, the expansion of GFP+ skeletal muscle mesenchymal progenitors can be detected on tissue sections at d3 post-fracture and polytrauma. By day 21 post-injury, skeletal muscle mesenchymal progenitors were increased around the fracture callus in polytrauma compared to fracture, coinciding with the abnormal accumulation of fibrotic tissue within the callus (Fig. 5a). Prx1 mesenchymal lineage gives rise to callus fibrosis that express the fibrotic markers Periostin and PDGFRα (Fig. 5b, c). Moreover, EDL and periosteum grafting showed that skeletal muscle mesenchymal progenitors produce bone and fibrous tissue in the callus, while periosteal cells only form bone by day 21 (Fig. 5d). To attenuate fibrosis, we treated mice with the clinically approved pan-tyrosine kinase inhibitor Imatinib^®^ that inhibits receptor phosphorylation including PDGFRα ^21, 22^. Imatinib^®^ treatment had no effect on bone repair after polytrauma by day 7. However, bone repair was improved as shown by decreased cartilage, bone and fibrosis volumes as compared to control by day 21 (Fig. 5e). Skeletal muscle surrounding the fracture site is therefore the origin of callus fibrosis and can be targeted pharmacologically to improve bone repair after musculoskeletal trauma.

**Figure 5:**
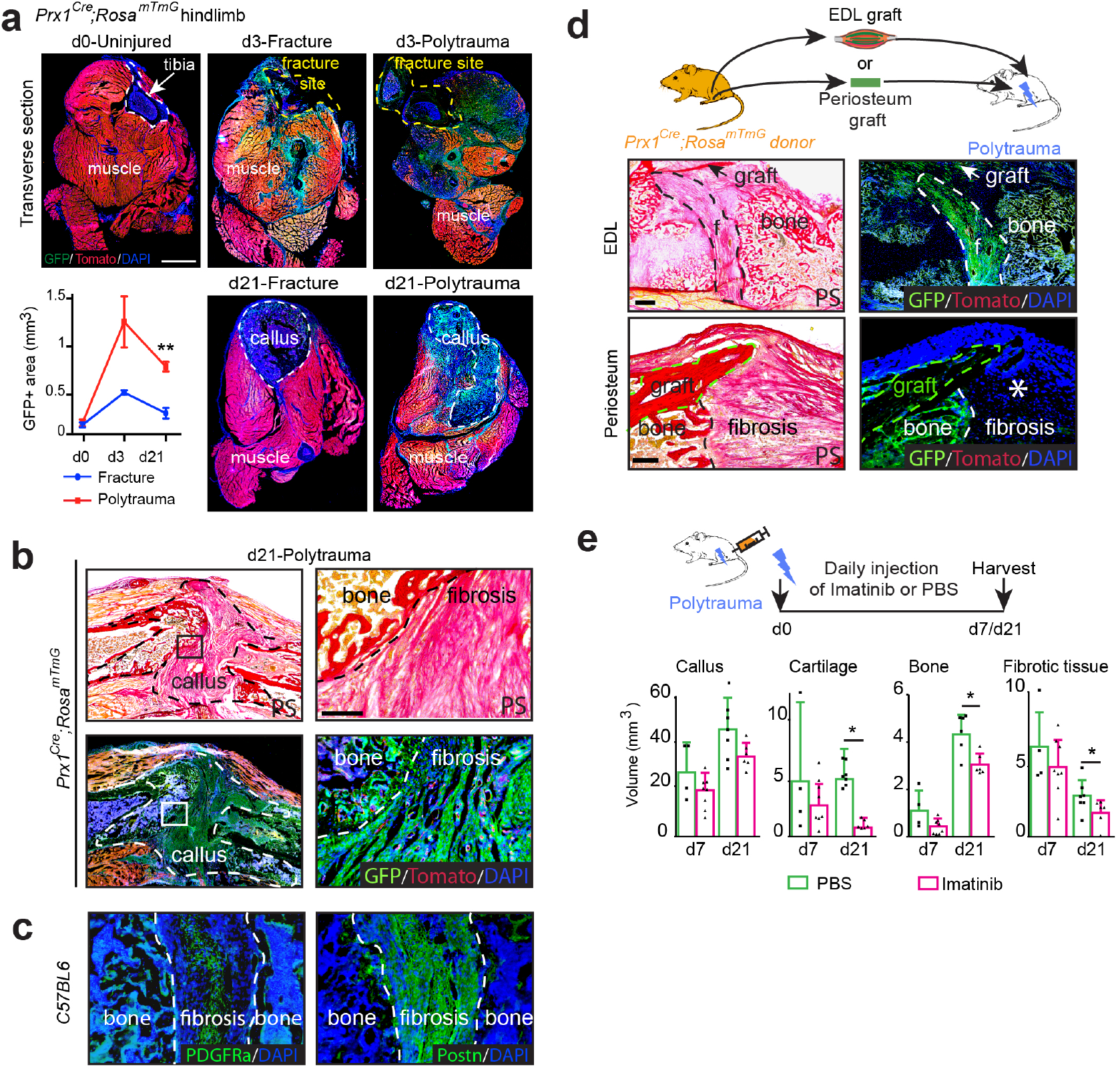
Skeletal muscle mesenchymal progenitors are the source of persistent callus fibrosis in polytrauma. **a**, Transverse sections of hindlimb from *Prx1^Cre^;Rosa^mTmG^* mice at day 0, day 3 or d21 post-fracture or post-polytrauma. Representative images of 3 distinct samples. Quantification of GFP+ area within skeletal muscle (callus excluded) on transverse sections of hindlimb of *Prx1^Cre^;Rosa^mTmG^mice* at d0 (uninjured), d3 and d21 post-fracture and post-polytrauma (d0 n=4, d3 post-fracture n=3, d3 post-polytrauma n=3, d21 post-fracture n=4 and d21 post-polytrauma n=3). T-test with Welch correction, **p<0.01 **b**, Longitudinal sections of fracture callus at day 21 post-polytrauma of *Prx1^Cre^;Rosa^mTmG^* mice stained with Picrosirius (PS, top) or counterstained with DAPI (bottom), and high magnification of boxed areas. **c**, Immunostaining of PDGFR*α* (green, left) and Periostin (green, right) in fibrosis of *wild-type* callus. **d**, Experimental design of EDL muscle or periosteum graft from *Prx1^Cre^;Rosa^mTmG^* mice transplanted at the fracture site in polytrauma (top). Longitudinal callus sections stained with PS and counterstained with DAPI at day 21 showed presence of GFP+ cells from EDL graft within bone and fibrosis and presence of GFP+ cells from periosteum graft within bone but absence within fibrosis (asterisk). **e**, Daily injection of Imatinib^®^ (50mg/kg/day) or vehicle (PBS) in mice with polytrauma. Histomorphometric analyses of total callus, cartilage, bone and fibrosis volumes of treated *vs* control mice at days 7 or 21 (d7 PBS-treated n=4, d7 Imatinib-treated n=8, d21 PBS-treated n=7, d21 Imatinib-treated n=6), two-paired Mann-Whitney test, \**P*=0.05. A, B, C, D, Representative images of 3 independent experiment. f: fibrosis. Scale bars: low magnification=1mm, high magnification=100μm. All data represent mean ± SD.

## Discussion

In this study, we uncover that skeletal regeneration implicates skeletal stem/progenitor cells from multiple tissue origins cooperating to repair bone. These skeletal stem/progenitor cells reside not only in bone (bone marrow, periosteum) but also in adjacent skeletal muscle and are all derived from a common Prx1-derived mesenchymal lineage. Previous work supported that cellular contribution of skeletal muscle to bone repair was restricted to specific conditions such as periosteal stripping and that non-skeletal mesenchymal stromal cells lacked chondro-osteogenic properties ^9, 12, 23^. Our results exclude the endogenous cellular contribution of the myogenic lineage during bone repair *in vivo* and demonstrate the direct contribution of skeletal muscle mesenchymal progenitors to cartilage and bone during bone repair. The skeletal muscle mesenchymal progenitors localize within skeletal muscle interstitium and overlap with FAP/MP population, highlighting news functions of FAP/MP as a heterogeneous and plastic population, which adapts its fate according to the environment^12, 24, 25^. Using scRNA-seq analysis, we show that within skeletal muscle mesenchymal progenitors, the FAP/MP population is distinct from pericytes, tenocyte-like and Spp1+/Lgals3+ populations and is the most responsive to bone injury. After fracture, we show that excepting Spp1/Lgals3+ population, FAP/MP, tenocyte-like cells, pericytes and fibroblasts adopt a fibrogenic state within 3 days post-injury associated with marked up-regulation of ECM genes. During this transient fibrogenic state, skeletal muscle mesenchymal progenitors express the master regulator of chondrogenesis Sox9 but are not engaged into a chondrogenic fate until day 5 post-fracture. Interestingly, lineage tracing showed that skeletal muscle mesenchymal progenitors are recruited at the fracture site between day 5 and day 7, indicating that they are already committed to chondrogenesis within muscle tissue before migrating at the center of the callus. Although skeletal muscle mesenchymal progenitors exhibit osteogenic potential *in vitro*, they preferentially engage in the chondrogenic lineage *in vivo* to support endochondral ossification.

In a polytrauma environment, the activation of skeletal muscle mesenchymal progenitors into the transient fibrogenic state is impaired and cells fail to undergo chondrogenesis. At later stages of repair, damaged skeletal muscle adjacent to the bone fracture is responsible for fibrous tissue accumulation within the facture callus, interfering with fracture consolidation. Skeletal muscle surrounding bone thus drives the fibrotic response and fibrotic remodelling during bone repair. Fibrosis is a dynamic process common to many tissue regeneration processes beginning with an initial phase of fibrotic response required for resident stem/progenitor cell activation and followed by active fibrotic remodelling necessary for completion of tissue regeneration ^26, 27^. In other regenerative processes, fibrotic progenitors are distinct from tissue-specific stem/progenitors and reside within the tissues themselves ^14, 28–30^. In bone regeneration, we showed that fibrotic progenitors are recruited from adjacent skeletal muscle after polytrauma and are derived from the same pool of progenitors that will undergo chondrogenesis. Several molecular therapies have been developed to treat fibrosis. Imatinib^®^, a pan-inhibitor of PDGFR, Bcr-abl and c-kit signalling pathways, ameliorates the late stages of bone repair in our polytrauma model suggesting potential applications of Imatinib^®^ or other related drugs in orthopaedics. Overall our findings have wide implications in musculoskeletal health, as they bring new knowledge on the role muscle plays during bone repair and define skeletal muscle mesenchymal progenitors as a prime target to enhance bone repair and prevents pathological fibrosis.

## Acknowledgements

We thank M. Garfa-Traoré, N. Goudin, C. Cordier, O. Duchamp de Lageneste, S. Alonso-Martin, B. Drayton, C. Masson for advice and/or technical assistance.

## Funding

This work was supported by INSERM ATIP-Avenir to C.C. and M.M, Fondation de l’Avenir to C.C., ANR-13-BSV1-001 to C.C. and F.R. and NIAMS R01 AR057344 and R01 AR072707 to C.C. and T. Miclau. A.J. was supported by a PhD fellowship from Paris Descartes University.

## Author contributions

A.J. performed the experiments with the help of A.K.. J.M. assisted with flow cytometry analyses. M.L. performed scRNAseq libraries. M.M. and F.R. reviewed the manuscript and gave advices. A.J. and C.C. designed the experiments, analyzed the data and wrote the manuscript. C.C. conceived the project and supervised the study.

## Competing interests

Authors declare no competing interests.

## METHODS

Further information and requests for resources and reagents should be directed to and will be fulfilled by Colnot Céline

### Mice

C57BL/6ScNj, *beta-actinGFP (GFP), Prx1^Cre^, Pax7^CreERT2^, Rosa-tdTomato-EGFP (Rosa^mTmG^*) and *Rosa^YFP^* mice were obtained from Jackson Laboratory (Bar Harbor, ME) and maintained on a C57BL6/J background. Mice were bred and kept under controlled pathogen conditions in separated ventilated cages in the animal facility of Imagine Institute, Paris. All procedures were approved by the Paris Descartes University Ethical Committee. Animals used for all experiments were males or females 10 to 14-week-old and experimental groups were homogeneous in terms of animal gender and age. No specific randomization methods were used. Sample labelling allowed blind analyses.

### Tamoxifen injection

To induce Cre recombination, Tamoxifen (TMX, T5648, Sigma) was prepared at a concentration of 10mg/mL diluted in corn oil, heated at 60°C for 2h and injected intraperitoneally (3 injections of 300μL per injection). *Pax7C^reERT2^* mice were injected with Tamoxifen at days 7, 6 and 5 before fracture.

### Tibial fractures and Polytrauma

For all surgical procedures, mice were anesthetized with an intraperitoneal injection of Ketamine (50mg/mL) and Metedomidine (1mg/kg) and received a subcutaneous injection of Buprenorphine (0.1mg/kg) for analgesia. The right leg was shaved and cleaned using Vetidine soap and solution (VET 001, Vetoquinol). For tibial non-stabilized fracture, the tibial surface was exposed, three holes were drilled through the cortex using a 0.4mm drill bit in the mid-diaphysis and the bone was cut to create the fracture via osteotomy. For polytrauma, the skin was incised and skeletal muscles surrounding the tibia were separated from the bone. Mechanical injury to skeletal muscles, including *tibialis anterior* (TA), *tibialis posterior, extensor digitorum longus* (EDL), *soleus, plantaris, gastrocnemius* muscles surrounding the tibia, was applied by compressing the muscles for five seconds along their entire length using a hemostat in a standardized and reproducible procedure. Tibial fracture was then created in the mid-diaphysis by osteotomy as described above. For skeletal muscle injury only, the same procedure was applied without bone fracture. For partial skeletal muscle injury, the same procedure was performed by compressing the TA and EDL skeletal muscles only prior to fracture. At the end of the procedure, skin was sutured using non-resorbable sutures (72-3318, Harvard Apparatus). Mice were revived with an intraperitoneal injection of atipamezole (1mg/mL), kept on heated plate and as soon as they were revived, they were allowed to ambulate freely. Mice received two supplementary doses of analgesia within 24h post-surgery and were monitored daily.

### Imatinib treatment

After polytrauma injury, mice received daily intraperitoneal injections of Imatinib^®^ (50mg/kg/day, Selleckchem, STI571) or vehicle (PBS) from the day of fracture until sacrifice.

### EDL skeletal muscle transplantation

Host mice received either a fracture or a polytrauma injury as described above. Donor mice were sacrificed by cervical dislocation and EDL skeletal muscle was dissected from tendon to tendon. The EDL skeletal muscle was transplanted adjacent to the fracture site at the time of fracture. EDL proximal tendon was sutured to the host patellar tendon and distal tendon was sutured to the host peroneus muscle tendon using non-resorbable sutures (FST, 12051-08). Skin was then sutured as described above. When fracture was induced one month after EDL muscle grafting, EDL was first grafted as described above along the tibia without fracture. One month later, after skin incision, the tibial fracture was performed as described above via osteotomy after carefully separating the grafted EDL muscle and the bone.

### Isolation of skeletal muscle cells and cell transplantation

*Prx1^Cre^;Rosa^mTmG^* mice were sacrificed by cervical dislocation. Skin and fascia were removed. *Tibialis anterior* (TA), *extensor digitus lengus* (EDL), *plantaris* and *soleus* muscles surrounding the tibia were dissected from tendon to tendon avoiding periosteum or bone marrow cell contamination. In a petri dish with 1mL of DMEM medium (21063029, Invitrogen), tendon and fat were removed and skeletal muscles were cut in small pieces using scissors. Skeletal muscles were transferred in 3mL of digesting medium containing DMEM (21063029, Invitrogen), 1% Trypsin (210234, Roche), 1% collagenase D (11088866001, Roche) and incubated in a water bath at 37°C for 2 hours. Every 20 min individualised cells were removed and transferred into ice-cold growth medium containing αMEM (32561029, Life Technologies) with 1% penicillin-streptomycin (P/S, 15140122, Life Technologies), 20% lot-selected non-heat-inactivated foetal bovine serum (FBS, 10270106, Life Technologies) and 10ng/ml bFGF (3139-FB-025/CF, R&D) and digesting medium was changed. This step was repeated until all skeletal muscle was digested. Cells were then filtered through 100μm filters (352360, Dutscher) and 40μm filters (352340, Dutscher) and centrifuged 10min at 1500 rpm and resuspended in appropriated volume of growth medium.

For cell sorting, skeletal muscle cells were resuspended in 1mL of sorting medium containing DMEM medium (21063029, Invitrogen) with 2% of FBS and 1% of P/S. Cell viability marker Sytox blue (1/1000, S34857, Thermofischer) was added just before sorting. Cell sorting was performed on BD FACS Aria II SORP (BD Biosciences) and cells were collected in growth medium. 150 000 freshly sorted skeletal muscle cells were embedded in TissuCol^®^ kit TISSEEL (human fibrogen 15mg/mL and thrombin 9mg/mL, Baxter, France) according to manufacturer’s instructions. Open fracture was performed as described above and cell pellets were transplanted at the fracture site.

### Callus sample processing, histology and histomorphometric analyses

Mice were sacrificed by cervical dislocation and fractured tibias were harvested at days 3, 5, 7, 14, 21, 28 or 56 post-surgery according to the experiment. Samples were fixed 24 hours in 4% PFA (15714, Euromedex) and decalcified in 19% EDTA (EU00084, Euromedex) for 21 days at 4°C upon agitation. Samples with endogenous expression of fluorescent reporters’ proteins were incubated in sucrose 30% at 4°C over night and then embedded in OCT (F/62550-1, MMFrance). All others samples were embedded in paraffin. All samples were sectioned throughout the entire callus and consecutive sections were collected. Tissue sections from paraffin embedded samples were incubated in NeoClear^®^ (1098435000, VWR) for 2×5min and rehydrated in successive baths of 100%, 90% and 70% ethanol and then rinsed in PBS for 5min. Frozen sections were let dry at room temperature for 30min and rehydrated in PBS for 10min. After staining, sections were dehydrated in 70%, 90%, 100% ethanol for 3min each and NeoClear^®^ for 10min. Slides were mounted using NeoMount^®^ mounting medium (1090160100, VWR).

#### Safranin’o staining

Nucleus were stained with Weigert’s solution of 5min. Slides were next rinsed with tap running water for 3min and then stained into 0.02% Fast Green for 30s (F7252, Sigma), followed by 30s into 1% acetic acid. To detect proteoglycan’s within cartilage, slides were stained with safranin’o solution for 45min (S2255, Sigma).

#### Masson’s trichrome staining

Sections were stained with Harris haematoxylin (dilution ½) for 5min (F/C0283, MMFrance), rinsed in running tap water 5min, then dyed with Mallory red for 10min, rinsed for 5min and then differentiated into phosphomolybdic acid 1% for 10min (HT153, Sigma). Collagen in bone was stained with light green for 20min (720-0335, VWR) and fixed into acetic acid 1%.

#### Picrosirius staining

Sections were stained in PicroSirus solution for 2h at room temperature, protected from light.

For histomorphometric analyses, every thirtieth section throughout the entire callus was stained with Safranin’o (SO), modified Massons’ Trichrome (TC), Picrosirius (PS) or counterstained with DAPI to visualize GFP signal and pictured using a Zeiss Imager D1 AX10 light microscope. Areas of callus, cartilage, bone, fibrosis and GFP signal were determined using ZEN software (Carl Zeiss Microscopy GmbH) and volumes were calculated via the following formula: 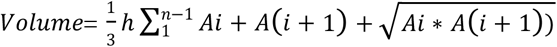 where Ai and Ai+1 were the areas of callus, cartilage, bone, fibrosis or GFP signal in sequential sections, h was the distance between Ai and Ai+1 and equal to 300 μm, n was the total number of sections analyzed in the sample.

### Skeletal muscle sample processing and analyses

*Tibialis anterior* (TA) skeletal muscle samples were harvested at specific time points, fixed for 3 hours in PFA 4%, incubated in sucrose 30% for 2 hours and embedded in OCT for cryosection. Sections were collected in the center of the muscle.

#### Haematoxylin-Eosin staining

Slides were stained with hemalun for 3min, rinsed in running tap water for 1min, stained with eosin for 1min (6766009, Thermo Fischer Scientific), then rinsed in water and pictured using a Zeiss Imager D1 AX10 light microscope.

### GFP quantification within skeletal muscle surrounding fracture callus

Tibial samples from *Prx1^Cre^;Rosa^mTmG^* mice were processed as described above for cryosection and were transversally included in OCT. Every fifth transverse section was collected in the middle of the diaphysis of uninjured tibia, at the fracture site day 3 post-fracture or in the center of the callus at day 21 post-fracture. Sections were counterstained with DAPI (eBiosciences, 495952) to allow GFP visualization. Entire transverse sections were pictured using Spinning Disk (Zeiss) and GFP signal was quantified within skeletal muscle excluding callus area using ZEN software (Carl Zeiss Microscopy GmbH).

### Immunofluorescence

GFP signal was detected without immunofluorescence staining. Cryosections were dried at room temperature for 30min, rehydrated in PBS for 10min and then mounted with Fluoromount (eBiosciences, 495952)^3^.

For calluses samples, paraffin embedded sections were deparaffinized and rehydrated as described above. For Periostin immunofluorescence, sections were blocked in 5% donkey serum for 1h and incubated in goat anti-periostin antibody (1/400, ref AF2955 R&D) diluted in 5% donkey serum in PBS (D9663, Sigma) over-night at 4°C. Sections were then washed in PBS 3×5min, incubated with donkey anti-goat AF488 antibody (1/250, A11055, Invitrogen) diluted in 5% donkey serum then rinsed with PBS for 10min and mounted using Fluoromount. For PDGFRα immunofluorescence, antigen retrieval were performed using sodium citrate buffer at 95°C for 20min. Slides were then cooled down in sodium citrate buffer on ice for 20min, rinsed in PBS 2×10min and then incubated in blocking solution containing 5% normal goat serum (NGS, Ab7481, Abcam), 0.5% Triton (T8787, Sigma) in PBS for 1h. Sections were washed in PBS 3×5min and incubated with goat anti-PDGFRα antibody (1/200, ref AF1062 R&D) diluted in blocking solution over-night at 4°C. Sections were then washed in PBS 3×5min, incubated with donkey anti-goat AF488 antibody (1/250, A11055, Invitrogen) diluted in blocking buffer then rinsed with PBS for 10min and mounted using Fluoromount. Samples were pictured using a Zeiss Imager D1 AX10 light microscope.

For skeletal muscle samples, cryosections were dried 30min at room temperature protected from light and then rehydrated for 10 min PBS. For anti-NG2, anti-PDGFRα and anti-PDGFRβ immunofluorescence, muscle cryosections were post-fixed in PFA 4% for 5min, rinsed 3×5min in 0.5% Triton in PBS, blocked in 5% NGS and 0.5% Triton diluted in PBS and incubated over night at 4°C with primary antibody: rabbit anti-NG2 (1/50, AB5320, Merck), goat anti-PDGFRα (1/200, AF1062, R&D) or rabbit anti-PDGFRβ (1/200, ab32570, Abcam). Slides were rinsed in PBS 3×5min and then incubated for 1h at room temperature in goat anti-rabbit AF647 (1/250, 21245, Life Technologies) or donkey anti-goat AF647 (1/500, ref ab150135 Abcam). Slides were mounted with Fluoromount (ref 495952, eBiosciences). For anti-CD29 immunofluorescence, cryosections were post-fixed in PFA 4% for 10min, washed for 2×5min in PBS, permeabilized in PBS-Triton 0.25%, blocked in 1% BSA for 15min (A2153, Sigma) and incubated with goat anti-mouse CD29 (1/50, 026202, R&D) overnight at 4°C. Slides were next rinsed and incubated in donkey anti-goat AF647 (1/500, ab150135, Abcam). Slides were mounted with Fluoromount. All pictures were obtained using a LSM700 confocal microscope (Zeiss).

For anti-αSMA immunocytofluorescence, cells were washed with PBS and fixed in PFA 4% for 15min, rinsed in PBS, permeabilized in PBS-Triton 0.25%, blocked in 5% NGS, incubated with anti-αSMA-Cy5 (ref AC12-0159-11, Clinisciences) for 1 hour and mounted with Fluoromount. Pictures were taken using a Zeiss Axio Vertical A1 light microscope.

### Flow cytometry

For flow cytometry analysis, skeletal muscle cells were incubated with CD31-PECy7 (PECAM-1, 561410, BD Biosciences), CD45-PECy7 (leukocyte common antigen, Ly-5, 552848, BD Biosciences), CD11b-PECy7 (integrin αM chain, 552850, BD Biosciences), CD34-AF700 (560518 BD Biosciences), Sca1-BV650 (740450, BD Biosciences), CD29-APC (130-096-306, Miltenyi Biotec), PDGFRα-BV711 (740740, BD Biosciences) and PDGFRβ-APC (17-1402-80, Invitrogen) for 15min on ice protected from light. Cells were then washed by adding 1mL of sorting medium and centrifuged for 10min at 1500rpm. Supernatant was trashed and cell pellets were resuspended in 200μL of sorting medium and Sytox blue was added just before analysis. Beads (01-2222-42, Thermo Fischer) were used for initial compensation set up and FMO controls were used to determine background level of each colour. Analyses were performed on BD LSR Fortessa SORP (BD Biosciences) and results analysed using FlowJo, LLC software, version 10.2. Gating strategy used for analyses is available in Extended Data Figure 4a.

### *In vitro* differentiation

After cell sorting, Prx1-derived skeletal muscle cells were expanded in 6-well plate to allow *in vitro* differentiation.

For adipogenic differentiation, sub confluent cells were placed into adipogenic medium containing αMEM supplemented with 10% FBS, 0.1μg/ml insulin (I3536, Sigma), 100μM indomethacin (I7378, Sigma), 0.5mM 3-isobutyl-1-methylxantine (I5879, Sigma) and 0.1μM dexamethasone (D8893, Sigma). Medium was changed every three days for two weeks. Lipid droplets were stained with Oil Red O solution (O0625, Sigma) and nucleus with Harris haematoxylin solution (F/C0283, MMFrance). For osteogenic differentiation, cells at confluence were cultured in osteogenic medium containing αMEM supplemented with 10% FBS, 0.1μM dexamethasone, 0.2mM L-ascorbic acid (A8960, Sigma) and 10mM glycerol 2-phosphate disodium salt hydrate (G9422, Sigma). Medium was changed every three days for three weeks. Mineralization was revealed with 0.2% alizarin red staining (A5533, Sigma). For chondrogenic differentiation, cells were plated as micromass at a concentration of 5.10^5^ cells in 200μL of growth media. Two hours later, growth medium was replaced by chondrogenic medium corresponding to DMEM with 10% FBS with 0.1μM dexamethasone, 100μg/mL sodium pyruvate (P5280, Sigma), 40μg/mL L-proline (P0380, Sigma), 50μg/mL L-ascorbic acid, 50mg/mL Insulin-Serine-Transferase (I1884, Sigma) and 10ng/mL TGFβ1 (T7039, Sigma). Medium was changed every three days for two weeks and proteoglycans were stained with alcian blue (A5268, Sigma). For myogenic differentiation, Prx1-derived muscle cells were plated at 1000 cells per cm^2^ and induced with myogenic medium containing F10 (31550-02, Life Technologies), 2% horse serum (26050088, Life Technologies) and 1% P/S for 3 days. For fibrogenic differentiation, cells were grown until sub-confluence and induced to fibrogenic differentiation with DMEM high-glucose (10566016, Life Technologies) with 10% FBS, 1% P/S and TGF-β1 at 1ng/mL. All pictures were obtained with a Leica DM IRB light microscope.

### RTqPCR analyses

Prx1-derived skeletal muscle cells at 80% of confluence were dissociated using trypsin, pelleted for 10 min at 1500 rpm and frozen at −80°C. RNA extraction was performed with RNAeasy Kit (74134, Qiagen) following manufacture’s instructions. Amount of RNA was quantified using NanoDrop 2000 UV-Vis Spectrophotometer (Thermo Scientific). 500μg of RNA were used to synthetize cDNA. RNAs were mixed with 1μL of oligo_12-18_ (18418-012, Life Technologies) and 1μL 10mMdNTP Mix (18427-013, Life Technologies) and heated at 65°C for 5min and left on ice for 1min. Next, 4μL 5X First-Strand buffer, 1μL 0.1M DTT, 1μL Superscript III RT^®^ (18080-044, Life Technologies) and 1μL RNaseOUT^®^ (10777-019, Life Technologies) were added and incubated at 50°C for 1h. The reaction was inactivated by heating at 70°C for 15min. qPCR mix was composed by 1μL of primers, 4μL of RNAse free H_2_O, 10μL of SYBR green Master Mix (11744-100, Life Technologies) and 5μL of cDNA. qPCR reaction was performed using 7300 Real-Time PCR System (Thermofischer Scientific). Mouse *Gapdh* was used as internal calibrator. qPCR analysis was done following ΔΔCT methods.

### Single cell analyses

Prx1-derived skeletal muscle cells were isolated as described above from *Prx1^Cre^;Rosa^mTmG^* d0 mice (un-injured), at day 3 post-fracture, day 3 post-polytrauma, day 5 post-fracture or day 5 post-polytrauma. Two mice were used per sample and only skeletal muscle was dissecting. Periosteum and bone marrow were not taken during the dissection. The scRNA-seq libraries were generated using Chromium Single Cell 3’Library & Gel Bead Kit v.2 (10x Genomics) according to the manufacturer’s protocol. Briefly, cells were counted, diluted at 1000 cells/μL in PBS+0,04% FBS and 20 000 cells were loaded in the 10x Chromium Controller to generate single-cell gel-beads in emulsion. After reverse transcription, gel-beads in emulsion were disrupted. Barcoded complementary DNA was isolated and amplified by PCR. Following fragmentation, end repair and A-tailing, sample indexes were added during index PCR. The purified libraries were sequenced on a HiSeq 2500 (Illumina) with 26 cycles of read 1, 8 cycles of i7 index and 98 cycles of read 2.

#### Aggregate sample generation

We generated aggregate sample to compared d0, d3 post-fracture, d5 post-fracture, d3 post-polytrauma and d5 post-polytrauma in a common dataset. Aggregate sample was generated according to cell ranger aggr pipeline in order to remove batch effect due to sequencing depth.

#### Seurat analysis

Seurat v3.1.2 and Rstudio v1.2.1335 were used for analysis of scRNA-seq data ^32, 33^. Cells expressing between 350 and 8000 genes and expressing less that 20% of mitochondrial gene were retained for analysis, genes expressed in less than 5 cells were not taken into account. Clustering was performed using the first 20 principal components with 0.5 as resolution and clusters were visualized using UMAP projection. Integrated analysis of d0, d3 and d5 post-fracture was performed using top 2000 features and the 20 first principal components with a resolution set at 0.5. Differentially expressed genes were determined using Wilcoxon rank sum test with P-value<0.05. For Gene Ontology (GO) analyses, differentially expressed genes were used to find enriched functions using Enrich R software (https://amp.pharm.mssm.edu/Enrichr/) ^34, 35^. GO functions including less than 5 genes and with adjusted adjusted P-value>0.05 were excluded. GO functions were classified into global functions and the percentage each function across all the functions found was plotted into radar graph as represented in the figure.

#### Monocle analysis

Monocle3 v0.2 was used for pseudotime analysis on FAP/MP of d5 post-fracture cells. Cells were ordered in a semi-supervised manner on the basis of Seurat clustering. Starting points correspond to the highest expression of *Ly6a* and *Cd34* genes.

#### Cell cycle analysis

Cell cycle analysis was performed using Cell Cycle Regression vignette from Seurat package.

#### Lineage analysis

Signature score was calculated for each cell as arithmetic mean of the expression of the associated genes in each cell (Table 1), and implemented as metadata in Seurat object.

### Statistical analyses

Data are presented as mean ± s.d. and were obtained from at least two independent experiments and n represents the number of samples used for the analysis. Statistical significance was determined with two-sided Mann-Whitney test and reported in GraphPad Prism v6.0a. Differences were considered to be significant when P <0.05.

## Notes

### Competing Interest Statement

The authors have declared no competing interest.

