## Supplemental Figures and Table for "Direct contribution of skeletal muscle mesenchymal progenitors to bone repair"

### Extended Data

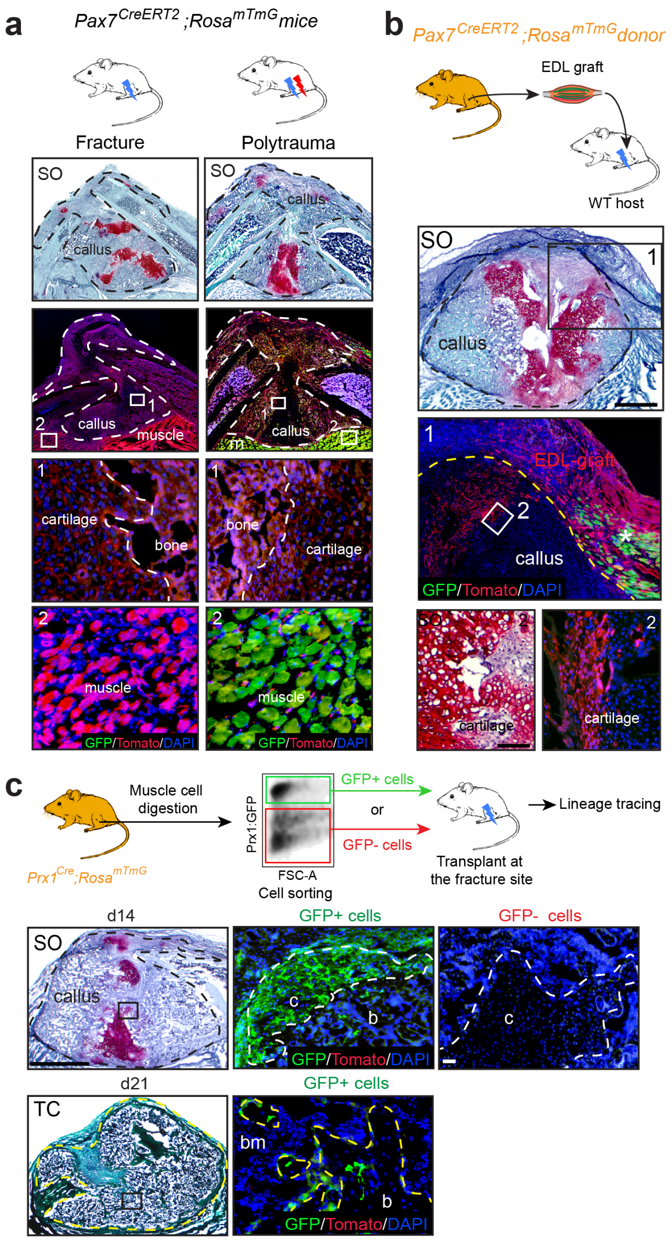


**Extended Data Fig. 1: Lineage analyses of skeletal muscle-derived cells contributing to bone repair,** related to Figure 1

**a**, Top: Experimental design. Bottom: Longitudinal callus section at d14 post-fracture (left) or polytrauma (right) of tamoxifen-induced *Pax7*$^{CreERT2}$*;Rosa*$^{mTmG}$ mice. High magnification shows no GFP signal in cartilage or bone (box 1) and GFP+ new myofibers next to callus in polytraumatic mice (box 2). **b**, Top: Experimental design of EDL muscle grafts from *Pax7*$^{CreERT2}$*;Rosa*$^{mTmG}$ mice transplanted adjacent to the fractured tibia of wild-type hosts. Bottom: Longitudinal callus section stained with SO at d14 post-fracture and adjacent section counterstained with DAPI showing EDL-derived cells (Tomato+) within the callus (delimited by a yellow dotted line) and regenerative myofibers within the graft (GFP+, asterisk). **c**, Experimental design of skeletal muscle cells isolated from skeletal muscle of *Prx1*$^{Cre}$*;Rosa*$^{mTmG}$ mice and transplanted at the fracture site of wild-type mice following cell sorting. Callus sections stained with safranin'o (SO) at d14 post-fracture and with Masson's Trichrome (TC) at d21 post-fracture and visualization of Prx1-derived and non Prx1-derived skeletal muscle cells on adjacent sections counterstained with DAPI. High magnification of cartilage (c, white dotted line) and bone (b, yellow dotted line) areas. Scale bar: SO/TC=1mm, high magnification= 50µm for cartilage and 25 µm for bone, bm: bone marrow. Representative images of 3 distinct samples.

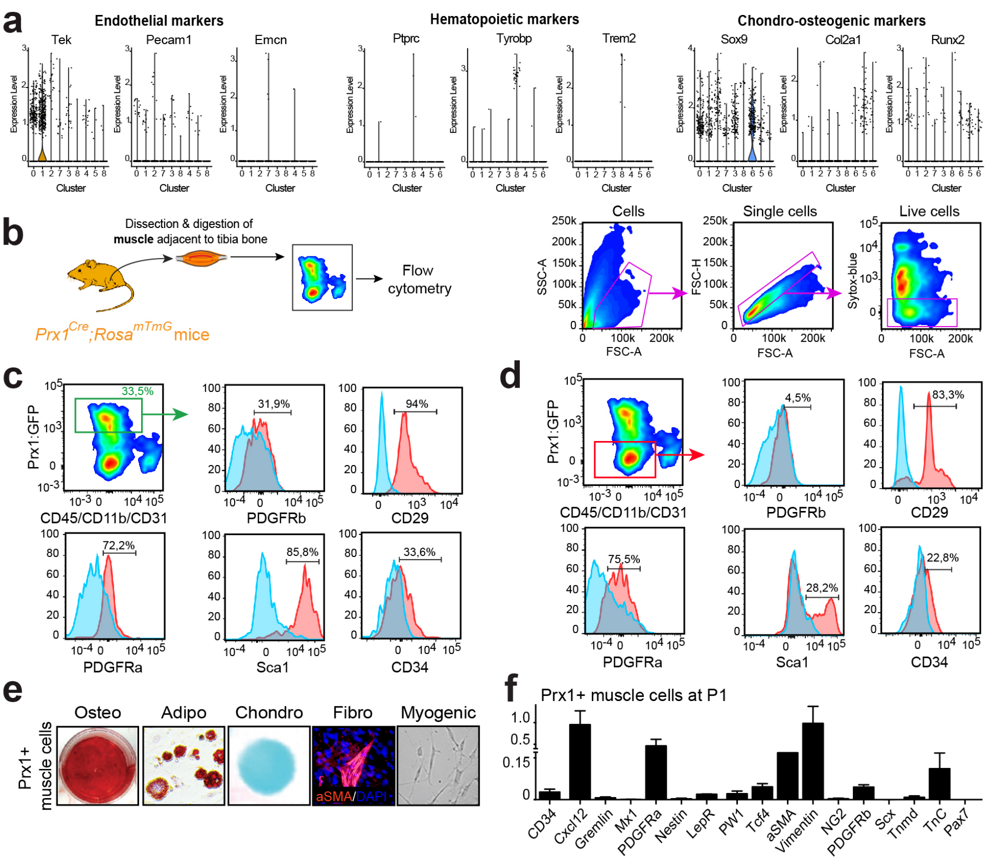


**Extended Data Fig. 2: Characterization of skeletal muscle mesenchymal progenitors,** related to Figure 2

**a**, Expression of endothelial, hematopoietic and chondro-osteogenic markers at d0 in scRNA-seq. **b**, Left, Experimental design of flow cytometry analysis. Right, Schematic representation of gating strategy of skeletal muscle cells used for flow cytometry analysis. **c**, **d**, FACS plots displaying the distribution of PDGFRβ, CD29, PDGFRα, Sca1 and CD34 cells in the GFP+/CD45-CD11b-CD31- population (**c**) and in the GFP/CD45-CD11b-CD31 double negative cell population (**d**). Red curve represents experimental tube and blue curve fluorescence minus one (FMO) control. Values represent the average of 3 independent experiments. **e**, Osteogenic (alizarin red staining), adipogenic (oil red o staining), chondrogenic (alcian blue staining), fibrogenic (aSMA immunocytochemistry) and myogenic (phase contrast) *in vitro* differentiation of Prx1-derived skeletal muscle cells. 3 independent samples were used to the analysis. Images are representative of 3 distinct experiments. **f**, RT-qPCR analysis on sorted Prx1-derived skeletal muscle cells from *Prx1*$^{Cre}$*;Rosa*$^{YFP}$ mice at P1. n=3 per group.

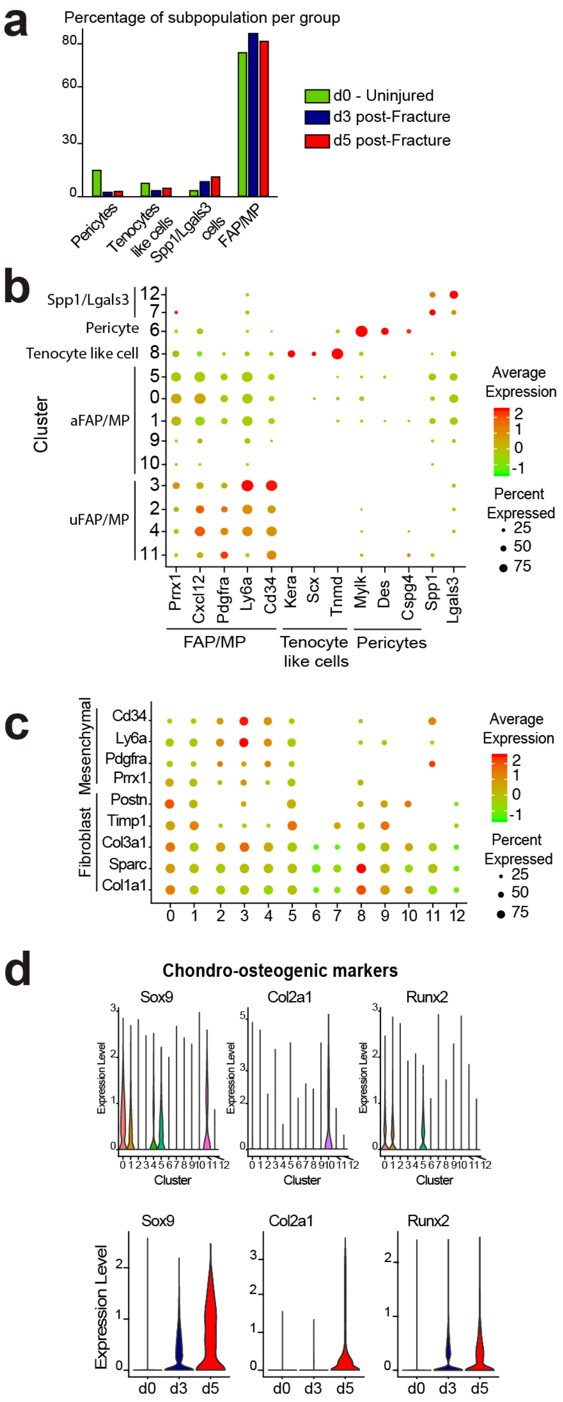


**Extended Data Fig. 3: scRNAseq analysis of skeletal muscle mesenchymal progenitors in response to fracture,** related to Figure 2

**a**, Percentage of subpopulation per sample in combined analysis of d0, d3 and d5 post-fracture samples. **b**, Dotplot of indicated genes expression identifying FAP/MP, tenocyte-like cells, pericytes and Spp1/Lgals3+ cell populations. **c**, Expression of fibroblast and mesenchymal markers per cluster. **d**, Expression of chondrogenic and osteogenic markers in d0, d3 and d5 post-fracture samples.

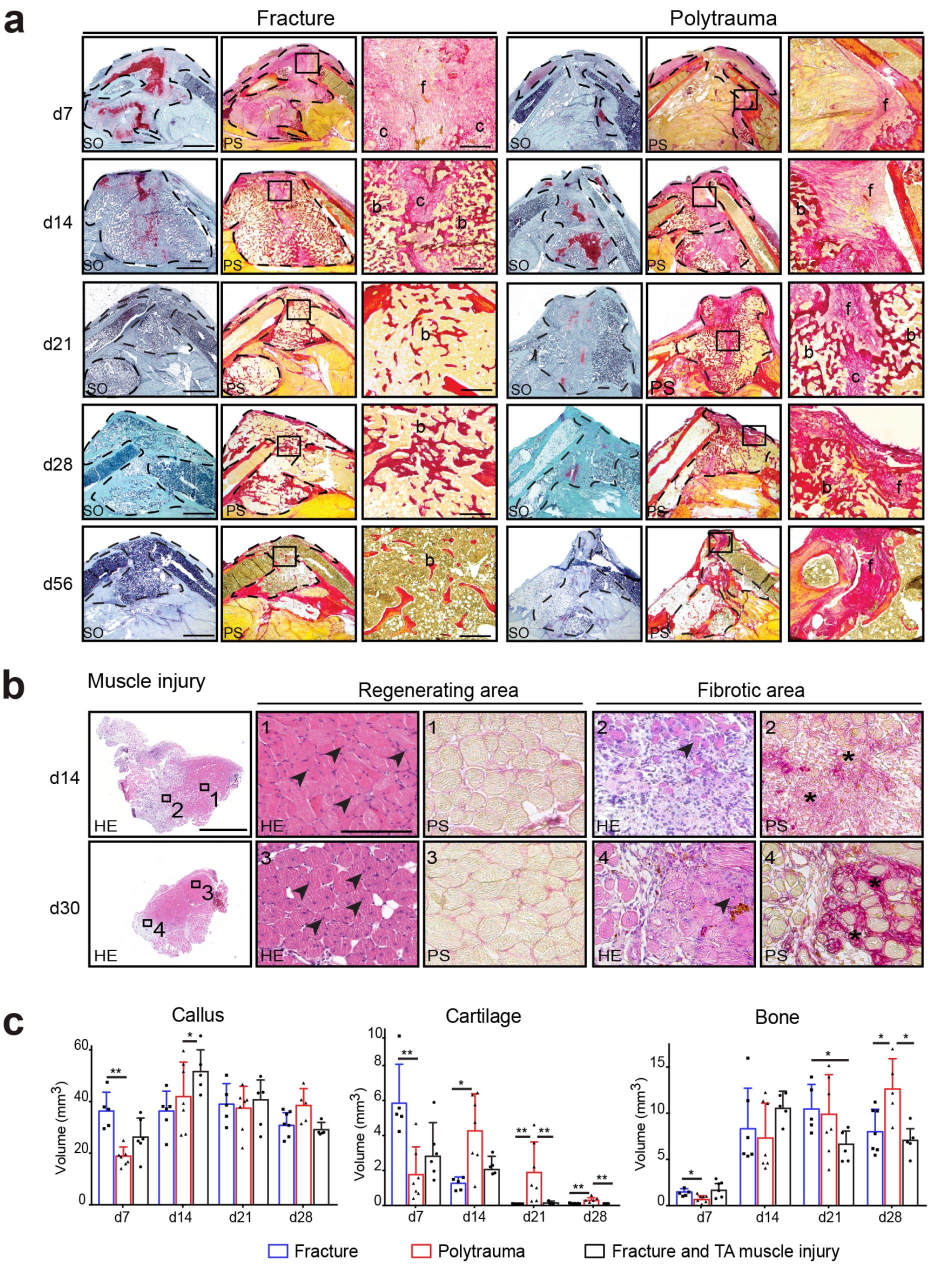


**Extended Data Fig. 4: Impact of skeletal muscle injury on bone repair,** related to Figure 3

**a,** Representative sections of fracture calluses stained with SO and PS at days 7, 14, 21, 28 and 56 post-fracture alone (left panels) and post-polytrauma (right panels) (callus outlined with a black dotted line, high magnifications of boxed areas). f: fibrosis, c: cartilage, b: bone, Scale bar=1mm, box areas =50µm. **b**, Tibialis anterior muscle sections at 14 and 30 days post-skeletal muscle injury alone (upper and lower panel respectively) stained with Hematoxylin and Eosin (HE) and Picrosirius (PS). High magnifications show centronucleated myofibers (arrowheads) in the regenerating area (boxes 1,3) and centronucleated myofibers surrounded by fibrous tissue (asterisks) in the fibrotic area (boxes 2,4). Representative images of 3 distinct experiments. **c**, Histomorphometric quantification of callus, cartilage and bone volumes in tibial fractures without muscle injury, with total muscle injury (as shown in Figure 1) or with TA muscle injury at days 7, 14, 21 and 28 post-fracture. d7 n=6, d14 n=5, d21 n=5, d28 n=5. Statistical analyses were performed following Mann-Whitney test, *$P$<0,05; **$P$<0,01, n=5 per group. All data represent mean ± SD.

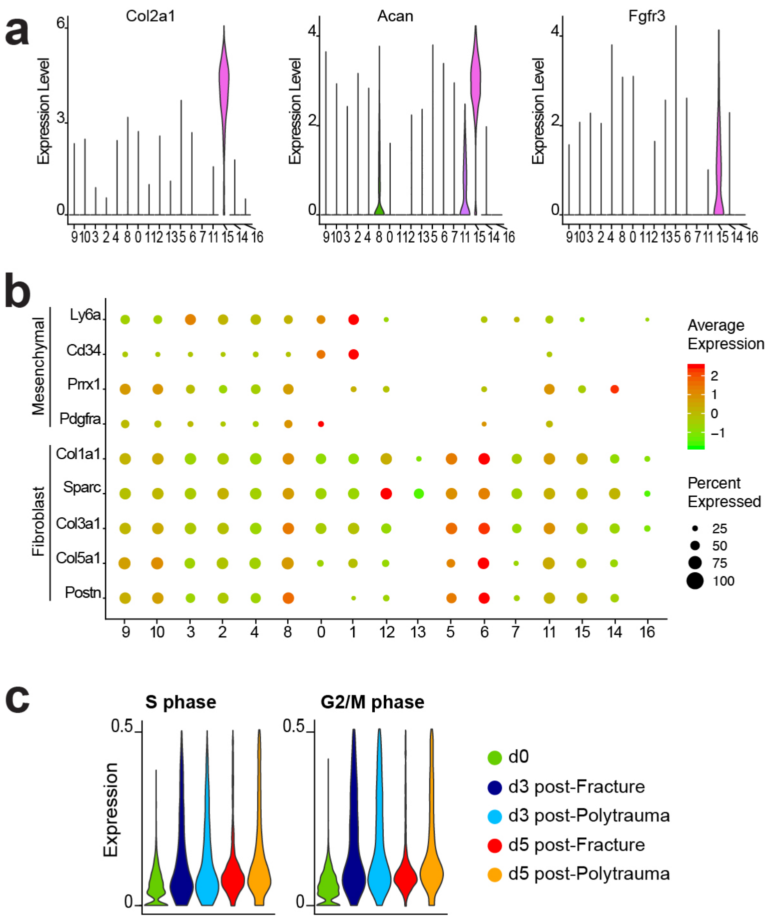


**Extended Data Fig. 5: scRNAseq analysis of skeletal muscle mesenchymal progenitors in response to fracture and polytrauma,** related to Figure 4

**a**, Expression of chondrogenic markers among clusters. **b**, Dotplot of fibroblast and mesenchymal markers among clusters in combined analysis of d0, d3 and d5 post-fracture and post-polytrauma samples. **d**, Expression of gene express in S or G2/M phases per sample.

| Lineage | Genes used |
|---|---|
| Fibrogenic | "Postn", "Aspn", "Tnn", "Tagln", "Col12a1", "Col5a1", "Sparc", "Col5a3", "Col6a2", "Col6a1", "Lum", "Col5a2", "Matn4", "Tnxb", "Col14a1", "Col16a1", "Col1a1", "Ogn", "Slc3a1", "Mmp19", "Adam12", "Thbs1", "Thbs2" |
| Mesenchymal | "Cd34", "Cxcl12", "Prrx1", "Ly6a", "Pdgfra", "Eng" |
| Osteogenic | "Alpl", "Sp7", "Ibsp", "Runx2" |
| Chondrogenic | "Col2a1", "Acan", "Sox9", "Pth1r", "Fgfr3", "Vegfa", "Wnt5a" |

**Extended Data Table 1: Markers used in lineage analysis,** related to Figures 2 and 4
